# A genome-based species taxonomy of the *Lactobacillus* Genus Complex

**DOI:** 10.1101/537084

**Authors:** Stijn Wittouck, Sander Wuyts, Conor J Meehan, Vera van Noort, Sarah Lebeer

**Affiliations:** Research Group Environmental Ecology and Applied Microbiology, Department of Bioscience Engineering, University of Antwerp, Antwerp, Belgium; Centre of Microbial and Plant Genetics, KU Leuven, Leuven, Belgium; Unit of Mycobacteriology, Department of Biomedical Sciences, Institute of Tropical Medicine, Antwerp, Belgium; BCCM/ITM Mycobacterial Culture Collection, Institute of Tropical Medicine, Antwerp, Belgium

## Abstract

**Background:** There are over 200 published species within the *Lactobacillus* Genus Complex (LGC), the majority of which have sequenced type strain genomes available. Although gold standard, genome-based species delimitation cutoffs are accepted by the community, they are seldom checked against currently available genome data. In addition, there are many species-level misclassification issues within the LGC. We constructed a *de novo* species taxonomy for the LGC based on 2,459 publicly available, decent-quality genomes and using a 94% core nucleotide identity threshold. We reconciled these *de novo* species with published species and subspecies names by (i) identifying genomes of type strains in our dataset and (ii) performing comparisons based on 16S rRNA sequence identity against type strains.

**Results:** We found that genomes within the LGC could be divided into 239 clusters (*de novo* species) that were discontinuous and exclusive. Comparison of these *de novo* species to published species lead to the identification of ten sets of published species that can be merged and one species that can be split. Further, we found at least eight genome clusters that constitute new species. Finally, we were able to accurately classify 98 unclassified genomes and reclassify 74 wrongly classified genomes.

**Conclusions:** The current state of LGC species taxonomy is largely consistent with genome data, but there are some inconsistencies as well as genome misclassifications. These inconsistencies should be resolved to evolve towards a meaningful taxonomy where species have a consistent size in terms of sequence divergence.

## Background

Since the advent of whole genome sequencing, it has become clear that current bacterial taxonomy is often not consistent with bacterial evolutionary history. In particular, many official taxa are polyphyletic and taxa of the same rank often differ in the diversity they represent in terms of sequence identity or shared gene content. The genus *Lactobacillus* is a prime example of both problems (Sun et al. 2015). First, it is larger than a typical bacterial family. Second, the genus is polyphyletic: the genera *Pediococcus, Leuconostoc, Weissella, Oenococcus* and *Fructobacillus* are also descendants of the most recent common ancestor of *Lactobacillus*. Together, these six genera form a monophyletic taxon that is often referred to as the *Lactobacillus* Genus Complex (LGC; Duar et al. 2017). The LGC is an important source of medical, food and feed applications (Sun et al. 2015) and is a key player in the human microbiome (Heeney, Gareau, and Marco 2018; Petrova et al. 2015). Because of all these reasons, the LGC taxonomy has been the topic of much study and debate, especially during recent years (Wittouck, Wuyts, and Lebeer 2019; Sun et al. 2015; Zheng et al. 2015; Salvetti et al. 2018).

In an attempt to systematically correct inconsistencies in the official bacterial taxonomy, Parks et al. (2018) recently constructed the Genome Taxonomy Database (GTDB). They built a phylogeny of dereplicated bacterial genomes across all phyla and determined the diversity within each official taxon using a heuristic called relative evolutionary divergence (RED). With this information, they corrected taxa that were polyphyletic or that had a diversity much above or below the average diversity of their rank. In their alternative, genome-based taxonomy, all six LGC genera were integrated into the family Lactobacillaceae, and the genus *Lactobacillus* was split into fifteen smaller genera.

The approach of Parks et al. to only split/merge taxa that are outliers in terms of diversity within their rank is appealing for the taxonomic ranks from phylum to genus, because these are relatively arbitrary and have no meaningful cutoffs in terms of diversity. The species level, however, is different. First, while there is agreement that the ranks from phylum to genus are arbitrary, there is discussion on the possible biological meanings of the species rank. For example, one recent study found that there exists a relatively large gap between within-species and between-species genome distances; in other words, that the “genome space” shows a discontinuity corresponding to the species level (Jain et al. 2018). In another recent study, the hypothesis was explored that taxa of the species rank show the property of exclusivity more than taxa of other ranks (Wright and Baum 2018). A taxon is exclusive if all of its members are more related to each other than to anything outside of the taxon. The authors concluded that their data falsified this hypothesis: many taxa, from many different ranks, indeed show the property of exclusivity, but that this was not especially often the case for taxa of the species rank. Thus, there is currently no agreement on whether a particular biological significance should be attached to the species rank. A second property of the species rank is more pragmatic, however. Contrary to the other ranks, relatively fixed similarity cutoffs to separate species are more or less generally accepted (Konstantinidis and Tiedje 2005; Jain et al. 2018).

The cutoff that is commonly used to separate bacterial species, based on their genome sequences, is 94-96% average nucleotide identity (ANI; Konstantinidis and Tiedje 2005; Richter and Rosselló-Móra 2009). Despite this cutoff being accepted as the new gold standard for bacterial species delimitation, it is seldom checked for type strain genomes of validly published species and subspecies. Type strain genomes of different species are sometimes more closely related than ∼95% ANI, while type genomes of subspecies of the same species are sometimes more distantly related than ∼95% ANI. For example, it has been shown that the type strains of *Lactobacillus casei* and *Lactobacillus zeae* are too closely related to consider them separate species (Judicial Commission of the International Committee on Systematics of Bacteria 2008). This has lead to the rejection of the *L. zeae* name. A related problem is that newly sequenced genomes, of type strains or otherwise, are not systematically checked for similarity against type strains. On the NCBI assembly database, uploaders of new assemblies are free to choose which species name they assign to the genomes. This has resulted in many misclassifications. For example, we have previously shown that many genomes annotated as *Lactobacillus casei* on NCBI are in reality more closely related to the *Lactobacillus paracasei* type strain (Wuyts et al. 2017). Similarly, strains of *Lactobacillus gallinarum* and *Lactobacillus helveticus* have often been wrongly classified (Jebava et al. 2014). Finally, it is likely that some sequenced genomes are so distantly related to all currently described species that they should be considered a new species.

In this work, we aimed to infer *de novo* species within the LGC by downloading all genomes belonging to this taxon from the NCBI assembly database and clustering them based on pairwise genome similarities. Advantages of this genome-based, *de novo* species taxonomy are that i) future work on comparing LGC species based on genomes can be done with a complete and non-redundant dataset of exactly one representative genome per species and ii) comparative genomics studies of individual species can proceed with all available, correctly classified genomes available for that species. Furthermore, we reconciled our *de novo* species with the currently known LGC species by identifying genomes of type strains and by comparing *16S rRNA* gene sequences to these of type strains. Based on this reconciliation, we propose changes to the official taxonomy in the form of splits or mergers of species.

As a genome similarity measure, we opted to compute the core nucleotide identities (CNIs) between all genomes. While ANI is the most commonly used genome similarity measure, it suffers from conceptual and practical drawbacks. First, since it is based on all genomic regions that are orthologous between two genomes, these regions will be different for each pairwise comparison. Thus, the type and amount of information are different across genome comparisons. Second, ANI is slow to compute for large datasets because it scales quadratically with the number of input genomes. The recently introduced fastANI (Jain et al. 2018) is orders of magnitude faster, but suffers from the same conceptual problem and remains an approximation. Assuming that core gene profiles or representative sequences are given, computing CNI similarities can theoretically be faster than ANI, because extracting the given core genes from all genomes scales linearly with the number of genomes. Using this principle, we developed a novel approach to rapidly obtain single-copy core genes (SCGs) for large genomic datasets. Briefly, we first identify candidate SCGs in a small, random subset of genomes and then extract those candidates from all genomes in the dataset using profile HMMs. Besides allowing for the calculating of CNI values, SCGs are also useful for two other purposes: genome quality control and tree inference. For quality control, they provide a biologically more meaningful alternative to the commonly used L50/N50 quality measures, by allowing an estimation of genome completeness and contamination (Parks et al. 2015; Waterhouse et al. 2017). Our pipeline for rapid SCG extraction was implemented in progenomics (a toolkit for prokaryotic comparative genomics) and is available at https://github.com/SWittouck/progenomics.

## Results

### SCGs and genome quality

We downloaded all 2,558 genomes that belong to the families Lactobacillaceae or Leuconostocaceae from genbank and identified single-copy core genes (SCGs) in this dataset. Our strategy for SCG extraction consisted of four steps: (1) full gene family clustering on a small, random subset of the genomes, (2) selection of candidate SCGs from these gene families using a very soft single-copy presence threshold, (3) search for the candidate SCGs in all genomes and (4) filtering of the definitive SCGs from the candidates using a fixed single-copy presence threshold. After visual inspection of the single-copy presence of the candidate SCGs in the full genome dataset (Figure S1), we retained candidate SCGs with > 95% single-copy presence. This strategy resulted in 411 SCGs *sensu lato* (“*sensu lato*” because we did not enforce 100% single-copy presence). We performed quality control of all genomes based on this set of SCGs (Figure 1). More specifically, we determined two quality measures for each genome: completeness, defined here as the percentage of *sensu lato* SCGs that are present in the genome, and redundancy, defined as the percentage of *sensu lato* SCGs showing two or more copies in the genome. The large majority of genomes had a completeness close to one and redundancy close to zero. Enforcing a minimum completeness of 90% and maximum redundancy of 10% for both measures resulted in 2,459 high quality genomes, or 96.1% of all genomes.

**Figure 1:**
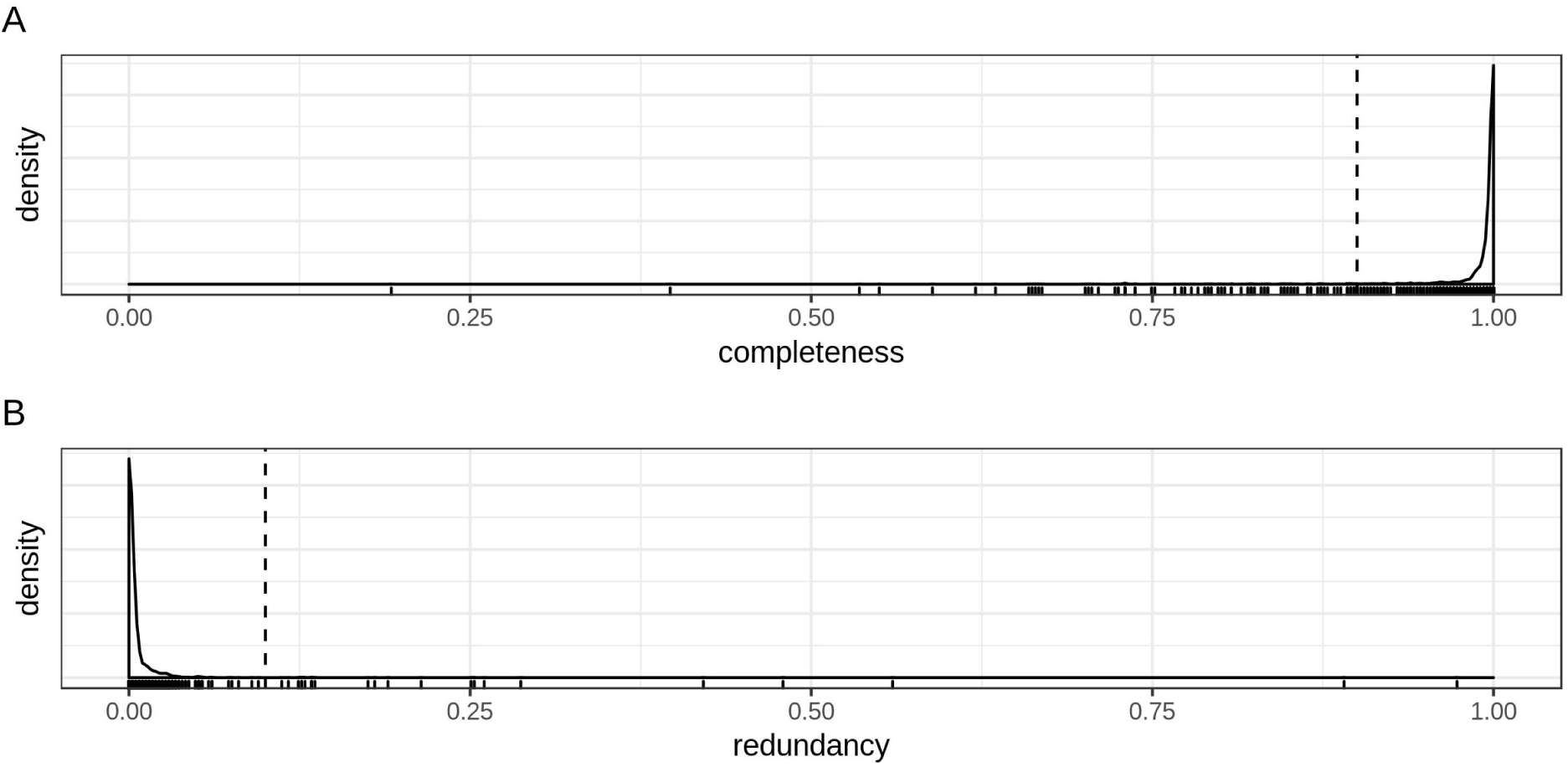
biologically meaningful genome quality control based on single-copy core gene (SCGs) completeness and redundancy. A) Density of genome completeness: for each genome, the percentage of SCGs that were present. B) Density of genome redundancy: for each genome, the percentage of SCGs with one or more extra copies. Small vertical bars at the bottom represent individual genomes and are shown to visualize outliers more clearly.

### Pairwise genome similarities and clustering

We compared every genome to every other genome using the core nucleotide identity (CNI), which we calculated from the supermatrix of SCGs. The density of these pairwise CNI values was strongly bimodal (Figure 2A): there was one sharp peak with CNI values > 94%, and a flatter peak (including multiple sub-peaks) with values < 90%. This indicated that two genomes were either very closely related (high CNI) or relatively distantly related (low CNI). Therefore, genomes were clustered into *de novo* species by grouping together genomes with similarities equal to or larger than 94%, using single linkage clustering. This resulted in 239 genome clusters.

**Figure 2:**
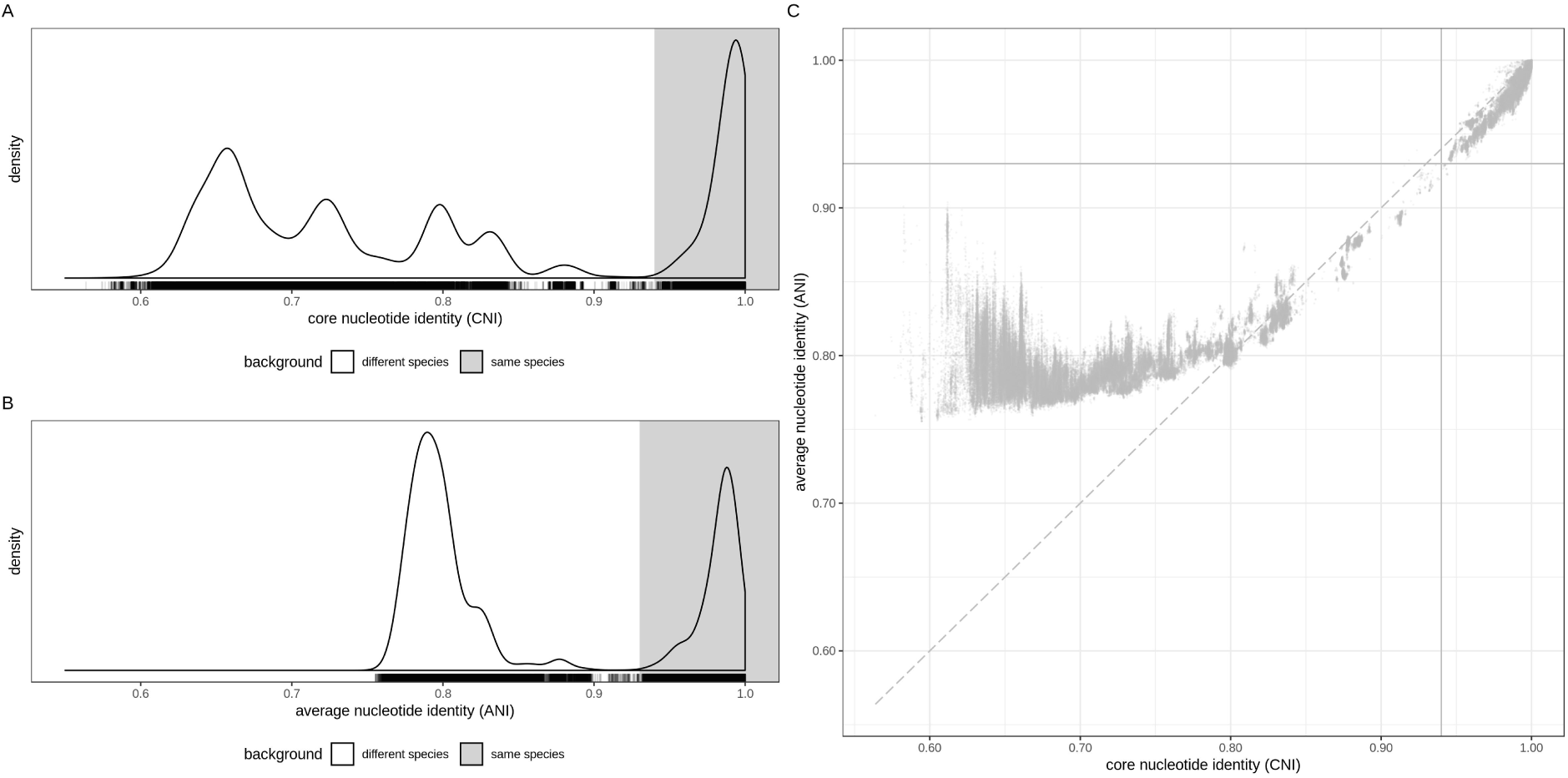
pairwise genome distance values. A) All pairwise core nucleotide identity (CNI) similarities. The grey area indicates the CNI values > 94% (the same-species range). B) All pairwise fastANI similarities. They grey area indicates the fastANI values > 93% (the hypothetical same-species range). C) CNI vs ANI values. Only genome pairs with fastANI > 75% are shown, since the fastANI tool doesn’t compute values under 75%.

One problem that can arise with single linkage clustering is that individual similarities within clusters can potentially get very small; much smaller than the clustering cutoff used. To assess whether this problem occurred for our clusters, we calculated for each cluster the minimum CNI between genomes within the cluster and the maximum CNI between genomes in the cluster and genomes outside of the cluster. Almost all within-cluster CNI values were larger than 94% (Figure S1). There was only one exception: two genomes in cluster 231 had a CNI of 93.6%, but even this value was very close to the species boundary. Furthermore, all 239 genome clusters met the criterion of exclusivity: all their within-cluster CNI values were larger than their between-cluster CNI values. This included cluster 231 (later identified as *Lactobacillus sake*i): its largest between-cluster CNI value was 81.6% (to cluster 221).

In addition to the CNI, we also calculated a second similarity measure between all our genomes: the fastANI similarity, which is a fast approximation of ANI. Just like the CNI distribution, the fastANI distribution was strongly bimodal (Figure 2B); its high-value peak covered the range 93% − 100%. A scatterplot of the CNI and fastANI similarities (Figure 2C) shows that both measures were strongly correlated for closely related genomes (around or above the species cutoff) and that the fastANI values were generally lower than the CNIs in this range. For more distantly related genomes, with CNI < 80%, the values were clearly less correlated, and the fastANI values were (much) larger than their CNI counterparts. The CNI species delimitation cutoff of 94% roughly corresponded to a fastANI cutoff of 93%. At these cutoffs, both similarity measures were in strong agreement on whether two genomes belonged to the same species or not.

To assess whether the species level held properties absent from the other taxonomic ranks, we calculated the exclusivity of clusters at various CNI cutoff values. The results showed that for CNI clustering cutoffs between 88% and 96%, all clusters were exclusive (Figure S2). This was not the case for most cutoffs outside of this range.

### Comparison with published species

We then attempted to label all 239 genomes clusters (*de novo* species) with the name of a published species. We did this in two stages: 1) identification of genomes belonging to type strains of published (sub)species (further called “type genomes”) and 2) for the remaining clusters: extraction of *16S rRNA* genes from the genomes and comparison to *16S rRNA* genes from type strains of published (sub)species (further called “type 16S genes”).

In the first cluster naming stage, we attempted to identify type genomes present in our dataset. First, we cross-referenced type strain names of validly published (sub)species found on the List of Prokaryotic Names with Standing in Nomenclature (LPSN; Euzéby 1997; Parte 2014) or Prokaryotic Nomenclature Up-to-date (PNU; DSMZ 2017) with the strain names that were listed in the NCBI assembly database. This resulted in the identification of 367 type genomes, belonging to 226 names (species or subspecies) from 211 species. Several type genomes for the same name were often found because many type strains have been sequenced multiple times. Next, we supplemented this type genome list by manually scanning the literature for extra type genomes, including also of species that were not published validly (i.e., not in an official taxonomic journal). This yielded type genomes for an additional 11 species. There were different reasons why manually identified type genomes had not been detected by our automated approach. Some of them were not represented on LPSN and PNU because they were not validly published. For other type genomes, their species were validly published, but too recently for them to be already on LPSN or PNU. *Lactobacillus nuruki* (Heo et al. 2018) was a validly published name but was not listed on LPSN or PNU, probably because its genus name was misspelled as “Lactobacilus” on NCBI and in the paper. *Convivina intestini* was missed by our automated approach because we did not include this genus in our search (while it does appear to be part of the LGC). The *Lactobacillus musae* type genome was missed because its type strain name (strain 313) consists only of numbers and was consequently filtered out by our pipeline. Finally, *Leuconostoc lactis* had a type genome on NCBI that did not pass our quality control, but we could also identify a type genome of *Leuconostoc argentinum*, a later synonym of *Leuc. lactis*. This type genome had not been identified by our automated pipeline because it filtered out rejected (sub)species names.

Of our set of 239 genome clusters, 210 contained one or more type genomes (Table S1). The large majority of these clusters (200 of them) contained type genomes from exactly one species (sometimes multiple type genomes from different subspecies or a resequenced type strain) and could thus be given the name of that species. Ten clusters contained type genomes from two or three different species, indicating that the type strains of these species were more closely related than the 94% CNI cutoff (Table 1). These clusters were: cluster 135 with *Lactobacillus amylotrophicus* and *Lactobacillus amylophilus*, cluster 141 with *Lactobacillus kimchii* and *Lactobacillus bobalius*, cluster 173 with *Lactobacillus micheneri* and *Lactobacillus timberlakei*, cluster 178 with *Weissella jogaejeotgali* and *Weissella thailandensis*, cluster 179 with *Lactobacillus fructivorans* and *Lactobacillus homohiochii*, cluster 218 with *Pediococcus acidilactici* and *Pediococcus lolii*, cluster 225 with *Leuconostoc gelidum, Leuconostoc inhae* and *Leuconostoc gasicomitatum*, cluster 233 with *Lactobacillus gasseri* and *Lactobacillus paragasseri*, cluster 236 with *Leuconostoc suionicum* and *Leuconostoc mesenteroides* and cluster 238 with *Lactobacillus casei* and *Lactobacillus zeae*. To these clusters, we assigned the name of the species that was earliest described. In one situation, two subspecies of the same species had a type genome in different genome clusters: *Lactobacillus aviarius* subsp. *aviarius* and *Lactobacillus aviarius* subsp. *araffinosus*. We named the genome cluster with the latter type strain *Lactobacillus araffinosus.*

**Table 1:**
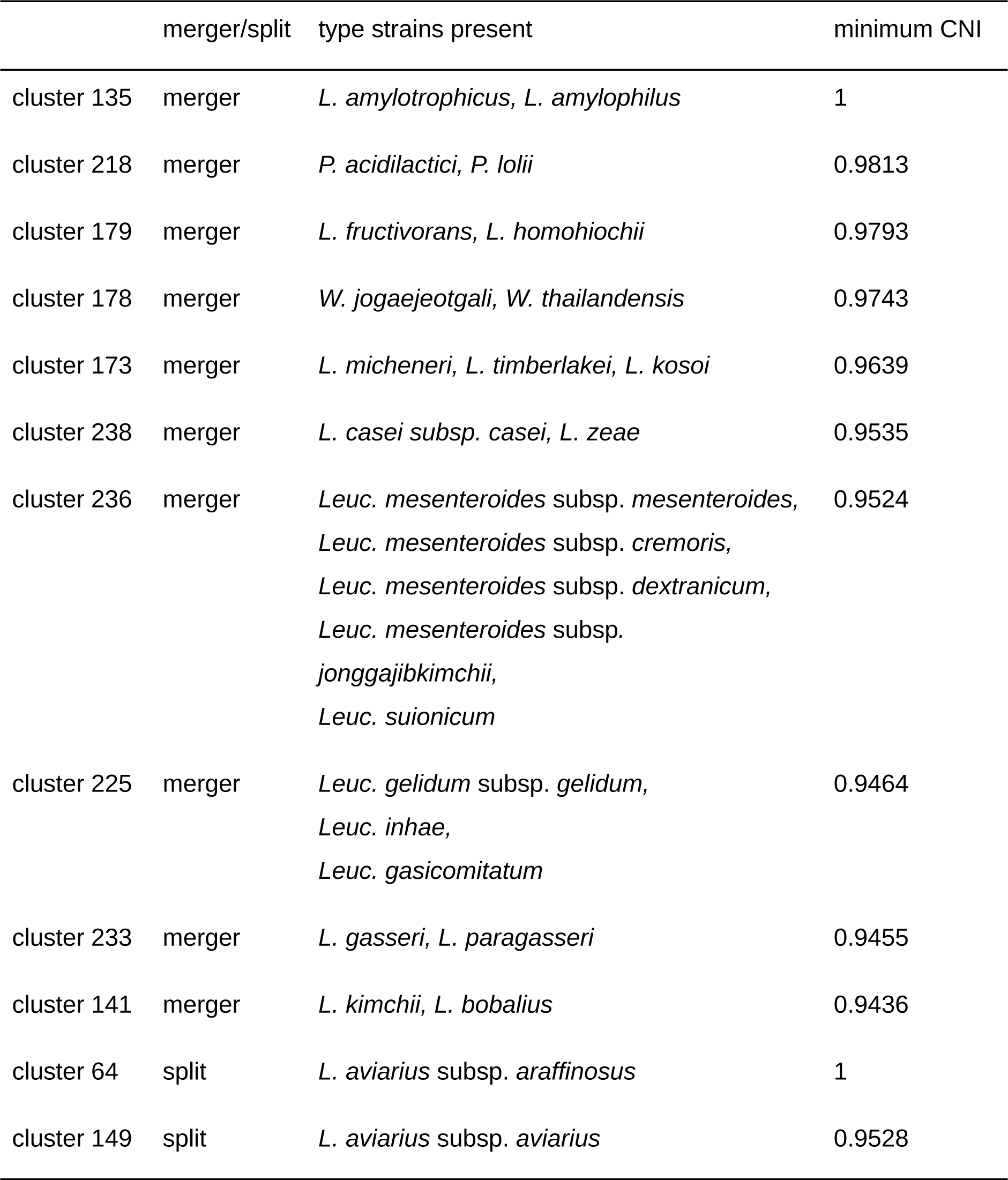
Inconsistencies between published and *de novo* species.

In the second cluster naming stage, we manually named the 29 remaining genome clusters in which we had not found type genomes (Table 2). We could assign a species name to twelve genome clusters with relative certainty because there was agreement between two sources of information: (i) the NCBI species labels attached to the genomes in the cluster and (ii) matches to our database of *16s rRNA* genes of validly published species not yet used to name a cluster via type genomes. For these genome clusters, we included a question mark in the species name to indicate that these names were not 100% certain, but rather a kind of “best guess”. Interestingly, we labeled eight clusters as “new species” because they yielded *16S rRNA* gene sequences that showed no hits to our type *16S rRNA* gene database, indicating that they were different from all known and validly published species. Each of these clusters consisted of exactly one genome. Finally, for the remaining nine clusters, the available information did not allow us to make a “best guess”, nor to consider them novel species with certainty. We labeled those clusters “unidentified species”.

**Table 2:**
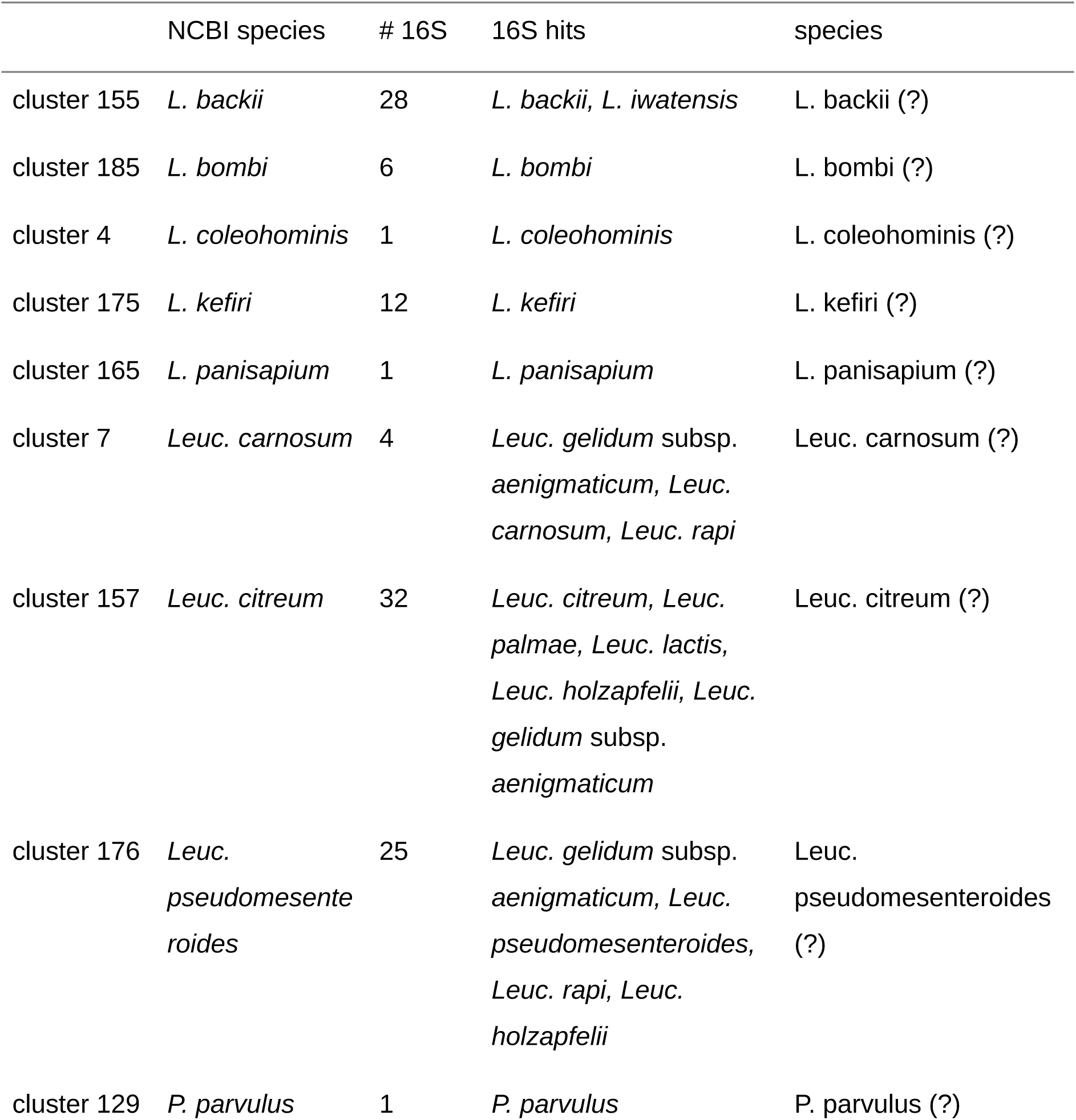
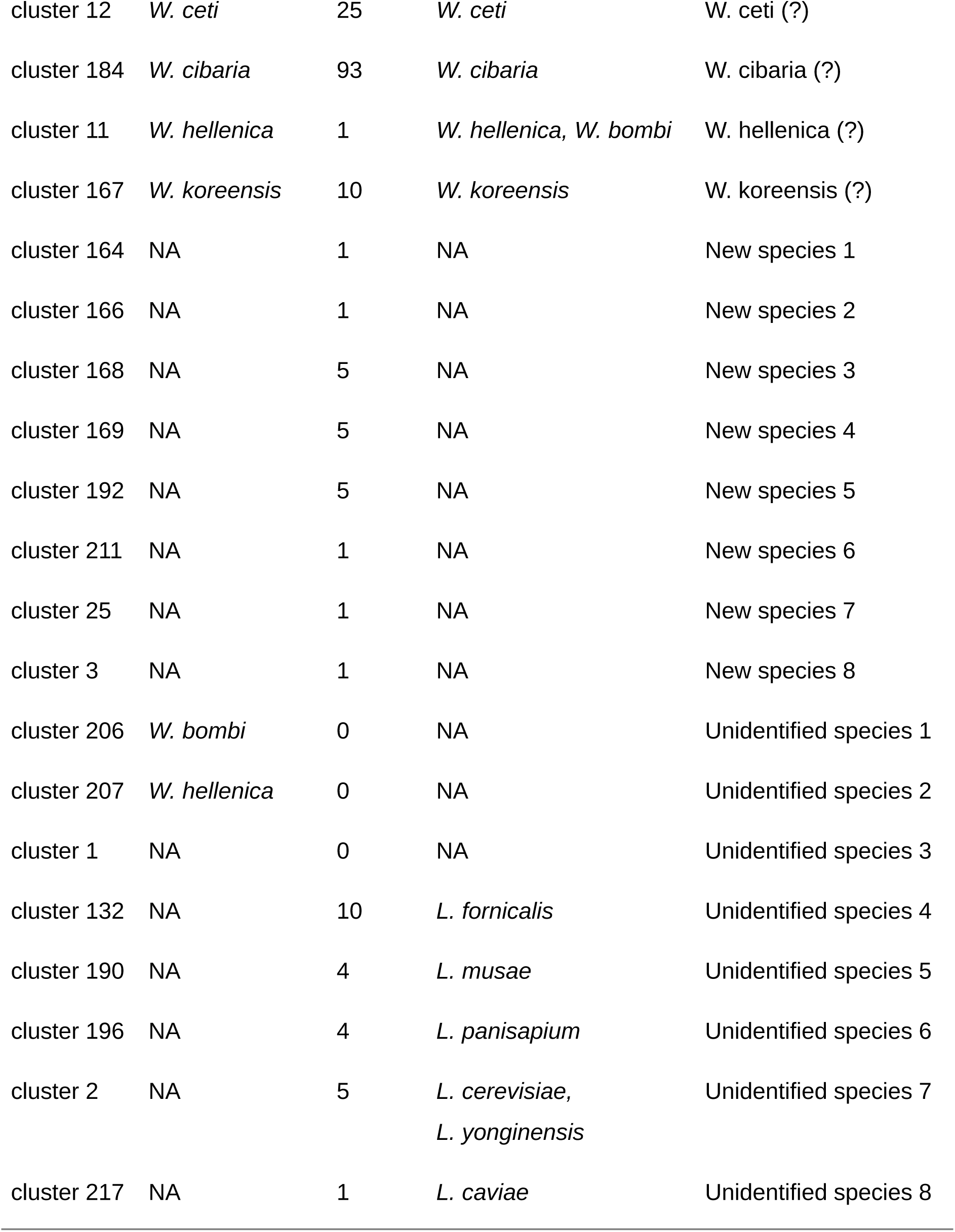
Manually named genome clusters.

We inferred a maximum likelihood species tree of the LGC using one representative genome (the best quality genome) for each of the 239 genome clusters (Figure 4). In this phylogeny, the eight new and nine unidentified species were also included. Two of the unidentified species were situated in the *Weissella* genus; the other new and unidentified species could be found in the *Lactobacillus* genus. Some of the new/unidentified species had long “tip branches” in the tree, indicating that they were relatively far removed from the other species. We annotated the tree with the phylogroups defined by Zheng et al. (Zheng et al. 2015) and predicted the lifestyle of the new species using the work of Duar et al. (Duar et al. 2017). Strikingly, the isolation source of each of the eight new species confirmed their predicted lifestyles, be it vertebrate-adapted, insect-adapted or free-living (Table 3). For nomadic phylogroups this was trivially the case, since they can occur in any environment.

**Table 3:** Phylogroups, predicted lifestyles and isolation sources of new species.

|  | phylogroup | predicted lifestyle | isolation source (NCBI) |
| --- | --- | --- | --- |
| New species 1 | L. delbrueckii group | vertebrate-adapted | a species of *Marmota* |
| New species 2 | L. reuteri group | vertebrate-adapted | a urine catheter |
| New species 3 | L. plantarum group | nomadic | kimchii |
| New species 4 | L. plantarum group | nomadic | kimchii |
| New species 5 | L. kunkeei group | insect-adapted | gut of *Bombus ignitus* |
| New species 6 | L. salivarius group | vertebrate-adapted | cow rumen |
| New species 7 | L. kunkeei group | insect-adapted | *Apis dorsata* |
| New species 8 | L. delbrueckii group | vertebrate-adapted | human gut |

**Figure 3:**
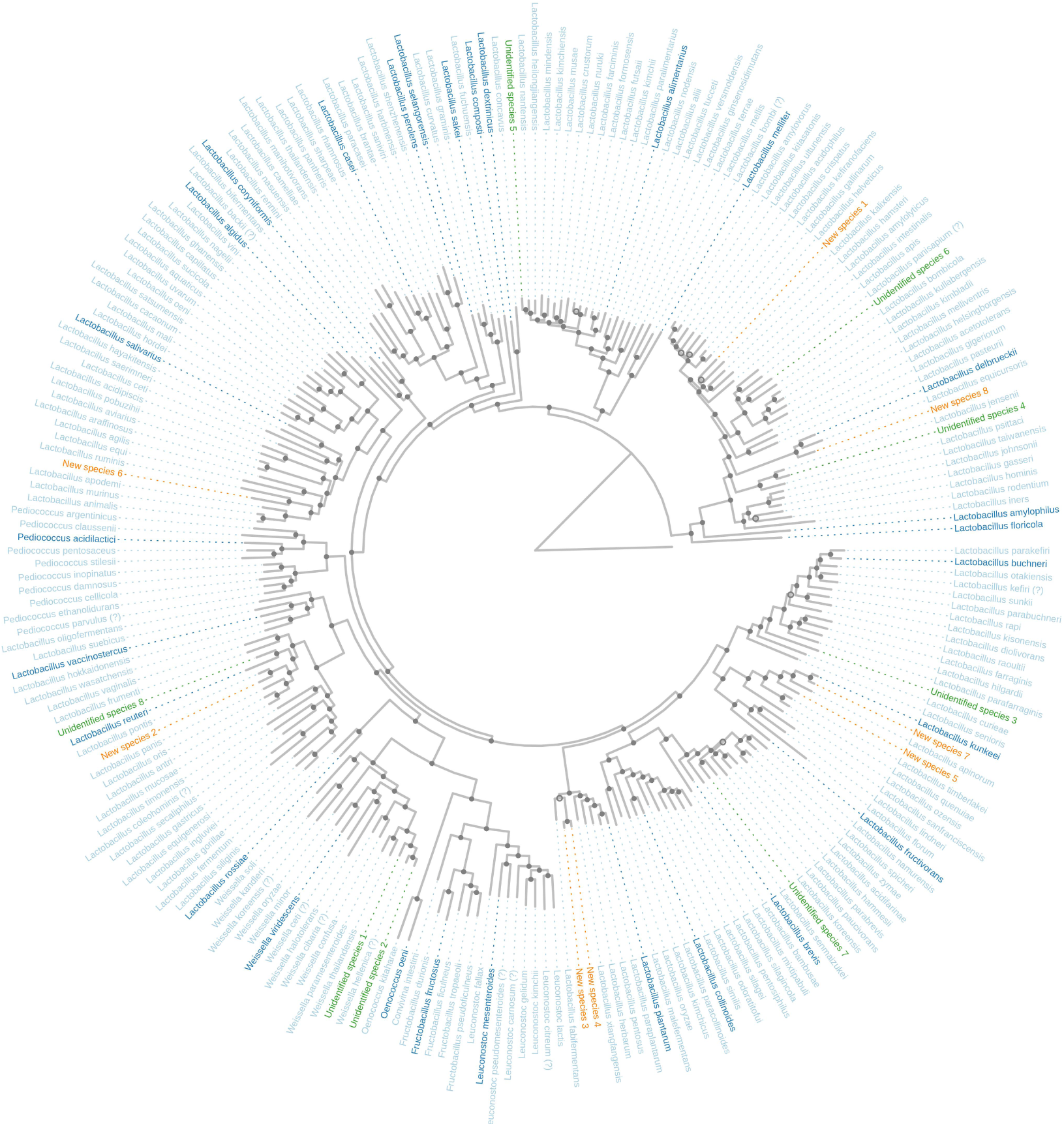
Maximum likelihood phylogenetic tree of all genome clusters. The tree was inferred on a nucleotide supermatrix of 100 SCGs and one representative genome per species. The genes and representative genomes were selected to maximize completeness of the supermatrix. The names and terminal branches of new and unidentified species are shown in orange and green, respectively. The names of “type species” of genera or phylogroups within *Lactobacillus* (Duar et al. 2017) are shown in darker blue. Weak clades (with a bootstrap value of < 70) are indicated with open circles. The root position was taken from the literature; the outgroup tip is artificial and its branch length was chosen in order to optimally visualize the tree topology.

**Figure 4:**
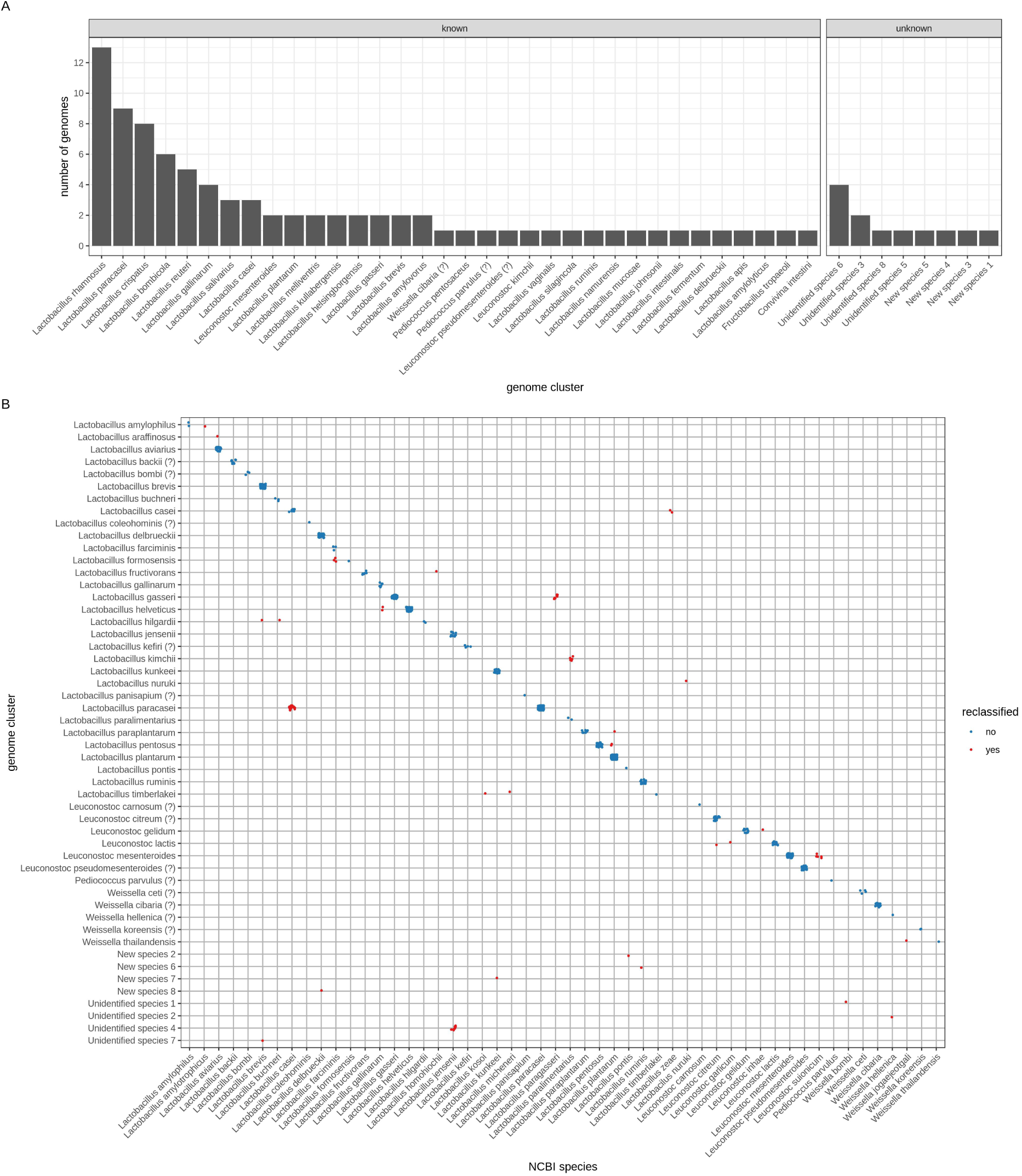
genome reclassifications. A) Classification of genomes that are currently unclassified on NCBI, using the CNI-based genome clusters. B) Reclassification of genomes that have an NCBI species label available but were found in a different CNI species cluster. Species names that are attached to an identical set of genomes in the NCBI and CNI classifications are not shown.

### Genome reclassifications

Of the 2,459 LGC genomes that passed quality control, 98 were unclassified at the species level on NCBI. As a direct result of our genome clustering and cluster naming pipelines, these genomes were automatically assigned to a species; either an existing one or a new or unidentified species (Figure 3A). The most frequently identified species in this group was *Lactobacillus rhamnosus*, with thirteen previously unclassified genomes assigned to it. Interestingly, several unclassified genomes belonged to clusters that we labeled as new or unidentified species.

Of the genomes that did have a species label on NCBI, 74 were reclassified to other species by our approach (Figure 3B). For twelve NCBI species labels, all of their genomes were reclassified to other species. For eleven of these, the reclassifications were consequences of the species mergers that we already discussed. The remaining case is that of *Leuconostoc garlicum*: all these genomes were reclassified to *Leuconostoc lactis*. Other reclassifications were clear cases of genomes that had been wrongly classified by the researchers that had uploaded them. We reclassified twenty genomes from *Lactobacillus casei* to *Lactobacillus paracasei*, five genomes from *Lactobacillus paralimentarius* to *Lactobacillus kimchii*, three genomes from *Lactobacillus farciminis* to *Lactobacillus formosensis*, two genomes from *Lactobacillus gallinarum* to *Lactobacillus helveticus*, two genomes from *Lactobacillus plantarum* to *Lactobacillus pentosus*, one genome from *Lactobacillus brevis* to *Lactobacillus hilgardii*, one genome from *Lactobacillus buchneri* to *Lactobacillus hilgardii*, one genome from *Lactobacillus plantarum* to *Lactobacillus paraplantarum* and one genome from *Leuconostoc citreum* to *Leuc. lactis*. Finally, some of the genomes with a species label on NCBI ended up in genome clusters considered by us as new or unidentified species.

## Discussion

We calculated pairwise CNI as well as fastANI distances between all high-quality genomes within the LGC. We observed that for relatively closely related genomes (CNI 80% − 100%), the fastANI similarities were smaller than the CNI similarities. This can be explained by the fact that the CNI is based exclusively on conserved genes, and these genes can be expected to mutate more slowly. This while the fastANI is based on all homologous genome regions between two genomes; these encompass also recently gained, potentially fast evolving genes. For more distantly related genomes (CNI < 80%), fastANIs were larger than CNIs. A possible explanation for this is that for these genome pairs, it becomes more difficult for an ANI tool to detect the homologous regions that show lower sequence identity; the fastANI distance might thus be based only on the homologous regions with higher sequence identity because they are easier to detect. Thus, the CNI might be a better reflection of evolutionary distance because it is based on a fixed set of (pre-aligned) genes.

The observed discontinuity in the CNI density we observed confirms observations made by others (e.g. Jain et al. 2018). This could be taken as evidence that the species level holds real biological meaning, apart from its usefulness as a taxonomic rank. Alternatively, one could argue that the discontinuity is a result of genome sampling bias. Indeed, there have been many comparative genomics studies lately where the aim was to compare as many strains as possible from the same species (e.g. Douillard et al. 2013; Martino et al. 2016), and this might have biased genome databases towards sets of closely related strains. However, the observation that only the CNI range of 88% to 96% shows full exclusivity of clusters is a stronger confirmation of the biological meaning of the species rank. This should be tested in more taxa to further explore whether the property of exclusivity is connected to the rank of species and to study the species question in general.

We clustered genomes of the LGC into *de novo* species using single linkage clustering with a threshold of 94% CNI, probably corresponding to approximately 93% fastANI. This cutoff value was lower than what is commonly suggested (Konstantinidis and Tiedje 2005; Richter and Rosselló-Móra 2009); it was chosen because it corresponded to the discontinuity we observed in the density of pairwise CNIs. A consequence of using a lower-than-usual species separation cutoff is that there can be some discussion on the species mergers that result from it, since type strains separated by relatively low similarity values could still end up in the same genome cluster. The other side of that coin is that the species splits we propose, as well as the new species, should be relatively undisputed: they are justified by strong separations between genomes.

By applying single linkage clustering with a threshold of 94% CNI, we implicitly defined a species as a set of strains for which it holds that each strain is at least 94% similar to at least one other strain in the species, and to not a single strain outside of the species. Using this species definition, we found that species membership in the LGC was almost transitive: all strains within a species were more than 94% similar to each other (there was only one exception to this). This has a number of interesting consequences. First, it means that, when using this species concept, species are very robust to strain/genome sampling; adding extra strains will not lead to mergers of species that were previously separated. Second, it means that for strain classification, we can calculate the similarity of the new strain to one (random) representative strain per species. When the strain is > 94% similar to one strain of a species, it will also be > 94% similar to all other strains in the species. Similarly, when the strain is < 94% similar to one strain of a species, it will also be < 94% similar to all other strains in the species. Finally, transitivity means that clustering with single linkage or complete linkage will yield identical results. Therefore, transitivity is a very useful property to have for a species definition and a set of species.

Four of the species mergers that we suggest are backed by very high CNI values between the species and are thus relatively “certain”. *Lactobacillus micheneri* and *Lactobacillus timberlakei* belonged to a cluster with a minimum CNI of 96.4. These species were described in the same recent publication (McFrederick, Vuong, and Rothman 2018), so it is not clear which name should be rejected in favor of the other. *Weissella thailandensis* (Tanasupawat et al. 2000) and *Weissella jogaejeotgali (Lee et al. 2015)* were present in a cluster with minimum CNI 97.4. In the *W. jogaejeotgali* paper, it was already mentioned that its type strain was closely related to *W. thailandensis*, with 99.39% 16S identity. *Lactobacillus fructivorans* (Charlton, Nelson, and Werkman 1934) and *Lactobacillus homohiochii* (Kitahara, Kaneko, and Goto 1957) were present in a cluster with minimum CNI 97.9. To our knowledge, it has not been suggested previously that these species could be merged. Finally, *Pediococcus acidilactici* (Lindner 1887) and *Pediococcus lolii* (Doi et al. 2009) were in the same cluster, with minimum CNI 98.1. It was already found by Wieme et al. that the *P. lolii* type strain DSM 19927 is a *P. acidilactici* strain (Wieme et al. 2012), so we can confirm their conclusion.

Some other cases of potential species mergers were less clear-cut. Before the species *Lactobacillus amylotrophicus* was introduced (Naser et al. 2006), with type strain DSM 20534, its type strain had been classified to *Lactobacillus amylophilus*, with type strain DSM 20533 (Nakamura 1981). We now found that these two type strain genomes are almost identical (CNI of ∼100%) at the level of their core gene sequences. However, the publication that introduced the species *L. amylotrophicus* (Naser et al. 2006) clearly shows that strain DSM 20534 is relatively different from strain DSM 20533, based on a comparison of pheS and *rpoA* gene sequences. Thus, we suspect that a mistake was made and that both genomes are actually the same strain, while the other type strain has not yet been sequenced. It is difficult to guess which of the type strains was actually sequenced (twice) but luckily, strain DSM 20533 was also sequenced by another institute. This second DSM 20533 genome was also identical to the other two. Therefore, it is highly likely that all three genomes are DSM 20533, the *L. amylophilus* type strain, and that the *L. amylotrophicus* type strain has not been sequenced yet.

In a 2012 paper (Pang et al. 2012), it was suggested that *Lactobacillus kimchii* (Yoon et al. 2000) and *Lactobacillus bobalius (Manes-Lazaro et al. 2008)* be considered as later heterotypic synonyms of *Lactobacillus paralimentarius* (Cai et al. 1999). In our results, the type strains of *L. kimchii* and *L. bobalius* were in the same cluster, with minimum CNI 94.4, while the *L. paralimentarius* type strain was present in a different cluster. The two clusters showed a maximum CNI value of only 93.9. Thus, we can confirm that *L. kimchii* and *L. bobalius* can be considered heterotypic synonyms of each other, and we do not object to also consider them heterotypic synonyms of *L. paralimentarius* since their clusters were extremely closely related. *Leuconostoc gelidum* (Shaw and Harding 1989), *Leuconostoc inhae* (Kim et al. 2003) and *Leuconostoc gasicomitatum* (Björkroth, Geisen, and Schillinger 2000) were present in the same cluster, with minimum CNI 94.6. The type strains of *Leuc. gasocimitatum* and *Leuc. inhae* even shared 99.5 CNI. Thus, the latter two species should definitely be heterotypic synonyms. To our knowledge, this has not been suggested before. These two type strains shared 94.8 and 94.6 CNI with the *Leuc. gelidum* type strain, respectively; just on the border of being the same species.

In 2018, three *Lactobacillus gasseri* (Lauer and Kandler 1980) strains were reclassified as *Lactobacillus paragasseri*, sp. nov. (Tanizawa et al. 2018). We found both species in the same cluster, with minimum CNI 94.6. Thus, this is a borderline case and we do not object to *Lactobacillus paragasseri* being considered a separate species. *Leuconostoc mesenteroides* (Tsenkovskil 1878) and *Leuconostoc suionicum* (Jeon et al. 2017) were in the same cluster, with minimum CNI 95.2. Leuc. suionicum used to be a subspecies of *Leuc. mesenteroides*, but it was recently suggested to upgrade it to species status (Jeon et al. 2017). Judging from our results, this seems acceptable but was maybe not necessary.

*Lactobacillus casei* (Orla-Jensen 1916) and *Lactobacillus zeae* (Kuznetsov 1959) were in the same cluster, with minimum CNI 95.35. There has been much confusion surrounding the names *L. casei, L. paracasei* and *L. zeae*. This is mainly the case because in 1971, strain ATCC 393 was chosen as the type strain for *L. casei* based on carbohydrate fermentation profiles (Hansen and Lessel 1971). In 1980, this type strain was included in the Approved Lists of Bacterial Names (Skerman, McGOWAN, and Sneath 1980). However, the strain later appeared to be relatively distant to the other historical *L. casei* strains. Subsequent proposals to change this illogical type strain (e.g. Dellaglio et al. 1991) were rejected because they violated the International Code of Nomenclature of Bacteria (Judicial Commission of the International Committee on Systematics of Bacteria 2008). Thus, strain ATCC 393 remained the *L. casei* type strain, which had some confusing consequences. First, the name *L. zeae* was rejected because its type strain was too closely related to strain ATCC 393. In addition, the name *L. paracasei*, whose type strain was very closely related to the historical *L. casei* strains, was kept alive. The consequence was that many of the strains that were historically classified to *L. casei* should now be classified to *L. paracasei*, and that all strain historically classified to *L. zeae* should now be classified to *L. casei* (Wuyts et al. 2017). In this work, we confirmed that the *L. casei* type strain and the (rejected) *L. zeae* type strain are closely related, but not so closely that it would be absurd to consider them different species.

The species *Leuconostoc garlicum* has one high-quality genome on NCBI, but it was reclassified to *Leuconostoc lactis* (Garvie 1960) *by our pipeline. The name Leuc. garlicum* occurs in a few publications but has never been published, validly or otherwise. Therefore, we suggest to not validate this species name and to classify these strains to the species *Leuc. lactis*. Finally, we found that *Lactobacillus aviarius* subsp. *aviarius* and *Lactobacillus aviarius* subsp. *araffinosus (Fujisawa et al. 1984)* were 91.07 CNI apart and therefore we suggest to rename the latter to *Lactobacillus araffinosus* comb. nov. To our knowledge, it was never suggested before that these should be separate species.

## Conclusions

We constructed a genome-based species taxonomy of the *Lactobacillus* Genus Complex by performing single-linkage clustering on 2,459 high-quality genomes with a 94% core nucleotide identity cutoff. We found that the resulting species were discontinuous, fully exclusive and almost fully transitive. Based on a comparison of the *de novo* species with published species, we proposed ten mergers of two or more species and one split of a species. Further, we discovered at least eight yet-to-be-named species that have not been published before, validly or otherwise. Finally, we have shown that our newly developed method to extract single-copy core genes allows for taxonomic studies on thousands of genomes on a desktop computer. We believe that correcting the published species in the direction of our *de novo* taxonomy will lead to more meaningful species that show a consistent diversity.

## Methods

### Downloading of genomes and gene prediction

All genome assemblies classified to the families Lactobacillaceae (NCBI taxonomy id 33958) or Leuconostocaceae (NCBI taxonomy id 81850) were downloaded from genbank using the script genbank_get_genomes_by_taxon.py that is part of the tool pyani, version 0.2.7 (Pritchard et al. 2015). Gene prediction was performed on all of these assemblies using Prodigal version 2.6.3 (Hyatt et al. 2010).

### Extraction of single-copy core genes (SCGs)

To be able to rapidly extract single-copy core genes (SCGs) from our large genome dataset, we devised a new approach based on seed genomes. Our strategy can be divided into four steps. In the first step, n seed genomes were selected completely at random and their genes were clustered into gene families. The software OrthoFinder (Emms and Kelly 2015) was used for gene family inference; it performs all-vs-all blastp (Altschul et al. 1990) search, normalizes the scores for sequence length and evolutionary distances between the genomes, applies a score cutoff per gene based on its best bidirectional hits and finally clusters the resulting weighted graph using MCL (Dongen 2000). This process resulted in gene families for the seed genomes only. In the second step, candidate SCGs were selected from all seed gene families by identifying gene families present in more than k seed genomes. The third step was the identification of these candidate SCGs in all genomes. For this purpose, the seed sequences for each candidate SCG were first aligned using MAFFT (Katoh et al. 2002). Next, profile HMMs were constructed from those alignments using the hmmbuild tool of HMMER (hmmer.org). All genomes were then scanned for occurrences of the candidate SCGs with the hmmsearch tool of HMMER. Finally, a homology score cutoff was determined for each profile HMM (see next paragraph for the details) and these cutoffs were applied to the HMMER hits to yield all “real” hits of the candidate SCGs in all genomes. In the fourth step, the final SCGs were determined by retaining only candidate SCGs with exactly one gene (copy) in at least p percent of all genomes.

For the training of profile-specific hmmer score cutoffs, the following strategy was used. First, if a gene was a hit of multiple profiles (multiple candidate SCGs), only the highest scoring profile was kept. Next, profile-specific cutoffs were determined as follows. Imagine we are selecting a score cutoff for one given profile. For each genome that has at least one gene among the hits of the profile, the best-scoring gene is considered a “true” hit. All other genes (second, third, etc best hits of genomes) are considered “false” hits. A score threshold is then determined that maximizes the F-measure of the hits; this is the harmonic mean of the precision and recall. In effect, this means that the threshold is set in such a way that as many as possible “best hits of genomes” are included in the results, while at the same time as many as possible results should be a “best hit of a genome”. The optimization was performed using the R package ROCR (Sing et al. 2005) This approach was applied separately to every profile HMM (each one representing a candidate SCG).

The SCG extraction strategy we described here was implemented in progenomics: a general, in-development toolkit for prokaryotic comparative genomics (github.com/SWittouck/progenomics). The parameter values used in this study were 30 for n (number of seed genomes), 25 for k (required number of seed genomes a candidate SCG should be present in) and 95 for p (required percentage of total genomes a final SCG should be present in). Software versions used were progenomics version 0.9, OrthoFinder version 2.1.2, blast version 2.6.0, MCL version 14-137, MAFFT version 7.407, HMMER version 3.1b2 and ROCR version 1.0.7.

### Genome quality control

For each genome, two quality measures were calculated based on the SCGs: the completeness (percentage of SCGs present in the genome) and redundancy (percentage of SCGs having more than one copy in the genome). Only genomes with > 90% completeness and < 10% redundancy were retained.

### Calculation of pairwise genome similarities

For every unique pair of genomes, two similarity measures were computed: the core nucleotide identity (CNI) and average nucleotide identity (ANI). For the CNIs, multiple alignment was first performed for each SCG on the amino acid level using MAFFT version 7.407 (Katoh et al. 2002) with the default parameters. Those amino acid alignments were then used as a reference to align the SCGs on the nucleotide level (reverse alignment). The aligned nucleotides were then concatenated into one large alignment (“supermatrix”). Next, pairwise nucleotide differences were calculated from this alignment using the distmat tool of EMBOSS version 6.6.0.0 (Rice, Longden, and Bleasby 2000), without correcting for multiple substitutions. Those were converted to similarities by subtracting them from 1. Pairwise ANIs were calculated using the tool fastANI (Jain et al. 2018), version 1.1, which computes a very fast and relatively accurate approximation of ANI.

### Clustering of genomes

Genomes were clustered into *de novo* species by joining together all genome pairs with a CNI similarity greater than 0.94. One could think of this as “nonhierarchical single-linkage clustering”. A simple algorithm to perform this clustering was implemented in R version 3.5.1 (R Core Team 2018).

### Identification of type genomes

To identify genomes of type strains (“type genomes”), we first downloaded assembly metadata for all of our genomes from NCBI, using the rentrez R package version 1.2.1 (Winter 2017). One of those metadata fields was the strain name of the assembled genome. Next, we compiled a list of all type strain names of validly published species of the six genera that make up the LGC. Since type strains are deposited in at least two but often many more culture collections, all of them have have at least two names. To collect those type strain names, we developed an R package called tidytypes (github.com/SWittouck/tidytypes, version 0.0.0.9000), which scrapes the websites LPSN (Euzéby 1997; Parte 2014), PNU (DSMZ 2017) and StrainInfo (Verslyppe et al. 2014). StrainInfo was only used to find extra synonyms of type strains of species and subspecies found on LPSN and PNU. By looking up those type strain names (including their many synonyms) in the assembly metadata from NCBI (which includes a strain name field), we were able to identify type genomes in our dataset on a large scale and in an automated manner. We then supplemented this list by manually identifying extra type genomes in the following way. We gathered a list of species names that were present in the NCBI assembly metadata, but for which we hadn’t already found a type genome. For each of those species, we looked up the type strain names in the original paper that described the species. We then inspected the assembly metadata of the genomes classified to that species on NCBI in an attempt to find type genomes that were missed by the automated approach (for example because the species were not validly published). The result of the automated and manual approaches was a complete list of type genomes in our genome dataset.

### Assignment of species names to genome clusters

Two different strategies were employed to assign species names to the genome clusters: one based on the type genomes present in the clusters and one based on comparisons to *16S rRNA* gene sequences of type strains. First, we used the list of type genomes to assign species names to genome clusters. If the cluster contained type genomes from exactly one species, we assigned this species name to the cluster. If it contained type genomes from more than one species, we concluded that those species were closely related enough to be considered heterotypic synonyms and assigned the oldest species name to the cluster. If two clusters contained type genomes from different subspecies of the same species, we proposed to upgrade the cluster with the non-type subspecies (e.g. *Lactobacillus casei* subsp. *tolerans*) to species status (e.g. *Lactobacillus tolerans*).

For the genome clusters that did not contain one or more type genomes, we predicted *16S rRNA* genes using barrnap version 0.9 (Seemann 2018). For all species and subspecies names on LPSN for which we did not find a type genome in our dataset, we downloaded a *16S rRNA* sequence from genbank. We then scanned the 16S sequences extracted from our genomes against this reference 16S database using the blastn tool of blast version 2.6.0 (Altschul et al. 1990). We then applied a percentage identity cutoff of 98% on those hits. When a genome cluster contained one or more genomes with a match to an official species through the 16S approach, and the same species was listed in the species labels of one or more genomes, we assigned this species name to the genome cluster as a “best guess”. When a genome cluster yielded 16S genes but none of them showed a match to the reference database, we considered it a new species. Finally, some genome clusters remained that could not be identified or be considered a new species; we labeled those “unidentified”.

### Inference of species phylogeny

For each genome cluster, the genome with the largest number of SCGs present was selected as a representative genome. We then selected the 100 SCGs that had the largest single-copy presence counts in those representative genomes. We then constructed a supermatrix with the selected genomes and SCGs, in the same way as described in the section *Calculation of pairwise genome similarities* (including reverse alignment). Next, we trimmed columns that had gaps in more than 1% of the genomes using trimal version 1.4.rev15 (Capella-Gutiérrez, Silla-Martínez, and Gabaldón 2009). We inferred a maximum likelihood tree using RAxML version 8.2.11 (Stamatakis 2014) using the general time reversible model for nucleotide substitutions and the CAT method for modeling mutation rate heterogeneity across columns. We used the “-f a” option of RAxML, which combines rapid bootstrapping (Stamatakis, Hoover, and Rougemont 2008) with a slow search of the tree space starting from the bootstrap trees. The number of bootstrap trees was determined by the “N autoMRE” option, which will stop looking for bootstrap trees when they don’t seem to add any extra information (Pattengale et al. 2009).

### Data processing and visualization

All processing and visualization of table-format data was done in R version 3.5.1 (R Core Team 2018) using the tidyverse set of packages, version 1.2.1 (Wickham 2017). Phylogenetic tree visualization and annotation was done using ggtree version 1.12.7 (Yu et al. 2016).

## Declarations

### Ethics approval and consent to participate

Not applicable

### Consent for publication

Not applicable

### Availability of data and material

NCBI assembly accession numbers for the genomes used in this study can be found in Table S1. The code used for this study was split over four repositories. The main pipeline is available at https://github.com/SWittouck/legen_pipeline. R Markdown scripts for the data analysis, including the creation of all figures and tables, is available at https://github.com/SWittouck/legen_data_analysis. This repository also contains the necessary data files and thus can be run on its own. Some of the steps of the pipeline, including the extraction of single-copy core genes, were implemented in the progenomics toolkit for prokaryotic comparative genomics for maximal reusability. This toolkit is available at https://github.com/SWittouck/progenomics. Finally, the scraping of taxonomic websites to obtain type strain names of published prokaryotic species was implemented in an R package called tidytypes, which can be found at https://github.com/SWittouck/tidytypes.

### Authors’ contributions

StWi and SL designed the study. StWi performed the analyses, while SaWu, CM and VVN provided important input on the bioinformatics. StWi drafted the manuscript. All authors provided feedback on the structure and writing of the manuscript. All authors read and approved the final manuscript.

## Acknowledgements

We would like to thank Camille Allonsius for checking the text for readability and clarity. In addition, we want to thank the entire Lab of Applied Microbiology at the University of Antwerp and the Computational Systems Biology group at the KULeuven for providing feedback on the general ideas and results. Finally, we want to acknowledge Rhambo (R.I.P.) and LAMBorGENEi for performing the computations.

