## Supplementary figures and images for "A genome-based species taxonomy of the *Lactobacillus* Genus Complex"

### figure S1

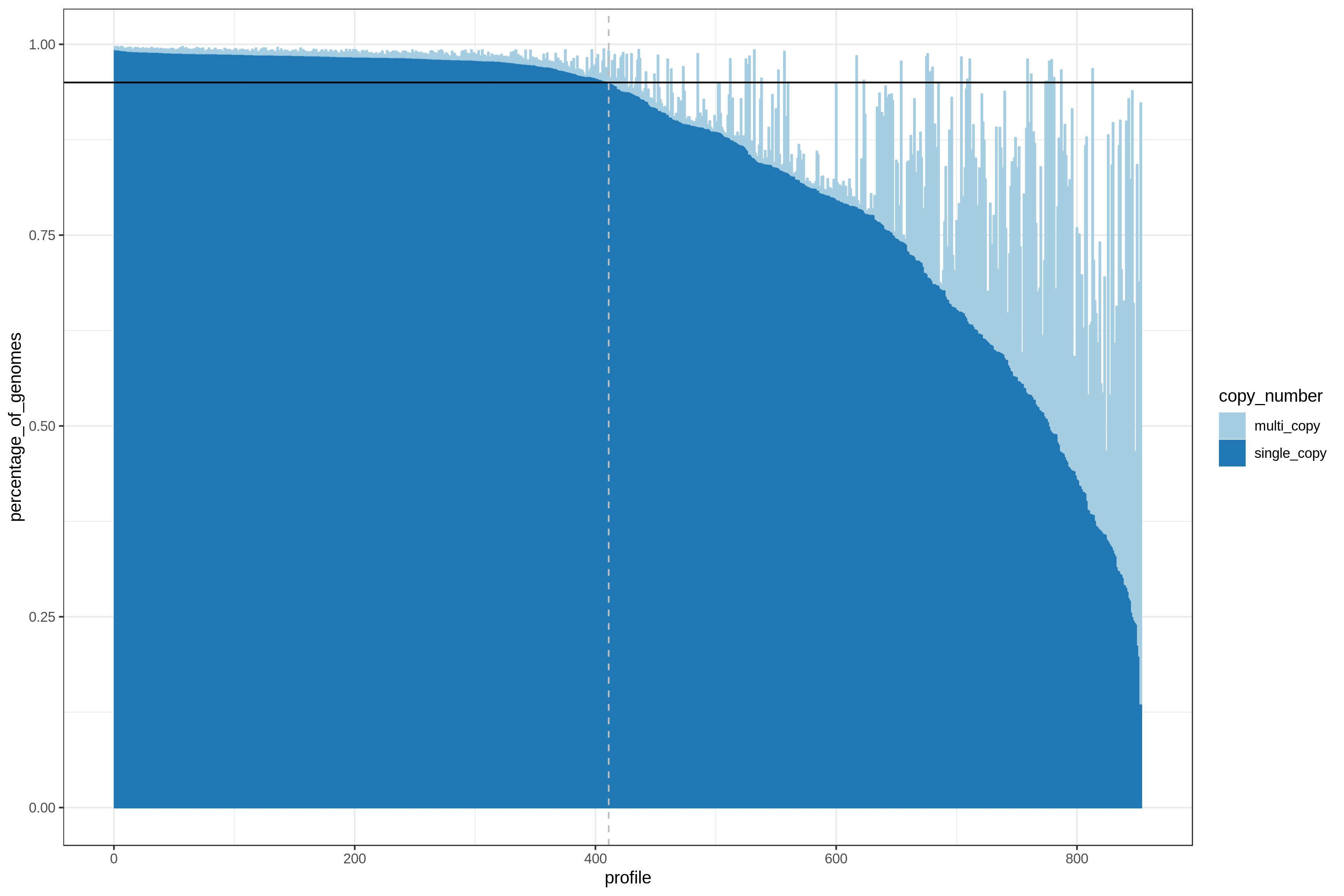

### figure S2

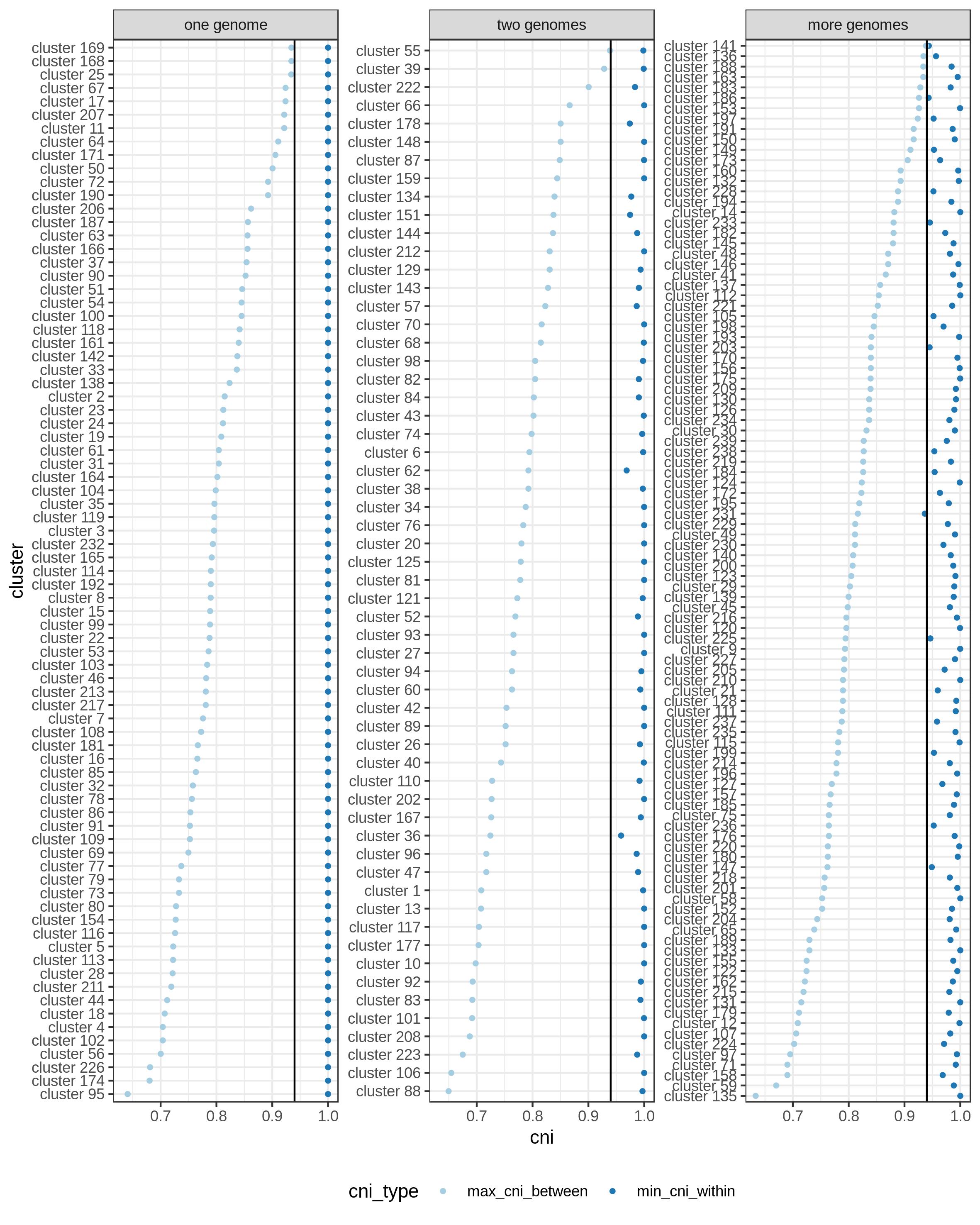

### figure S3

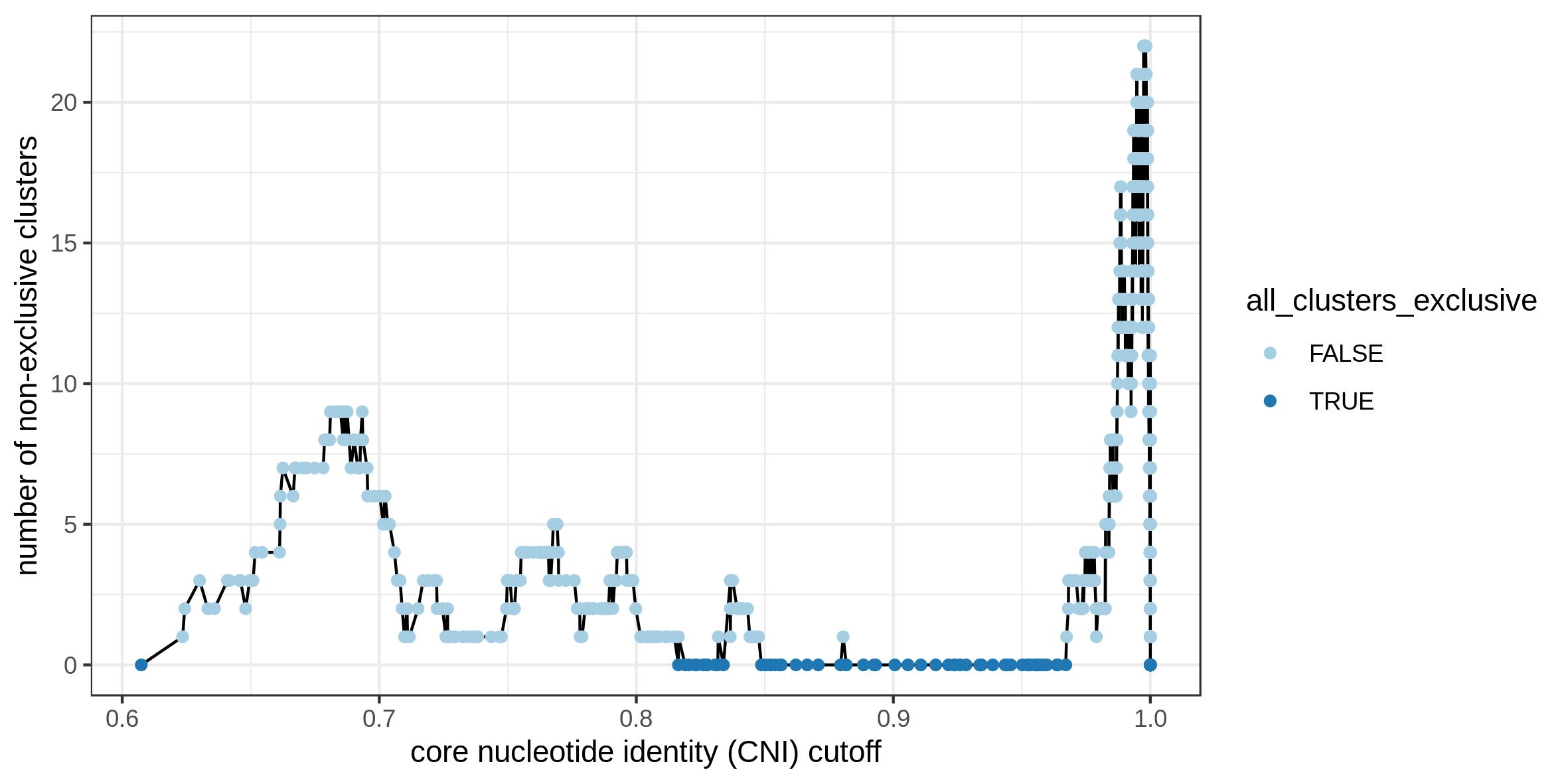
